# An epigenetic change in a moth is generated by temperature and transmitted to many subsequent generations mediated by RNA

**DOI:** 10.1101/099259

**Authors:** Jaroslav Pavelka, Simona Poláková, Věra Pavelková

## Abstract

Based on the presented results, the epigenetic phenomenon (paramutation) found in the short antennae (*sa*) mutation of the flour moth (*Ephestia kuehniella*) is probably determined by a small RNA (maybe piRNA) and transmitted in this way to subsequent generations through the male and female gametes. It is triggered by a change in the ambient environment; it persists for many generations (20 so far); during the epigenetic effect the original wild type appeared again. The epigenetic effect of the flour moth is induced by changes in ontogenetic development, such as increased temperature on pupae development, little nutritious food, different salts in food of certain chemicals into eggs. We found this mechanism may explain the intermittent clearance of this effect at some individuals and/or progeny of a pair in the generation chain in which the effect transfers. As the nature of the observed phenomenon could related to sirtuin genes, we hypothesize an association of these genes and non-coding small Piwi-interacting RNAs (piRNAs). The key is that the survival of RNA over many generations carries a number of practical implications. It is evident that the reaction to environmental change can manifest through RNA. It follows that there may be evolutionary significance in the long-term transmission of traits to future generations.

## Introduction

Paramutations are deviations from Mendelian inheritance. An important part of this phenomena, called an *epigenetic inheritance*, seems to be due to heritable alterations without a change in the sequence of the DNA. Epigenetic information is frequently erased at the start of each new generation. However, in some cases epigenetic information can be transmitted from parent to progeny (multigenerational epigenetic inheritance) (Buckley et al., 2012).

In our older study (Pavelka and Koudelová, 2001), we described a specific phenomenon of phenotypic inheritance (paramutation), which can be studied in the *short antennae* (*sa*) morphological mutation of the Mediterranean flour moth [*Ephestia kuehniella* Zeller (Lepidoptera: Pyralidae)]. In the case of the *sa* paramutation, the transmission to subsequent generations occurs through the male, as well as the female. Moreover the phenomenon reported here has other aspects: 1) it is triggered by a change in the ambient environment (e.g., temperature, LiCl in food); and 2) under certain conditions it rarely returns to the original state. The *sa* mutation is inherited as a simple autosomal recessive gene and causes considerable shortening of antennae in moths of both sexes. At higher temperature the antennae of *sa* moths revert to the wild-type (genotype remains the same), which is then transmitted to subsequent generations (Pavelka and Koudelová, 2001).

In summary, the short antennae (*sa*) high temperature mutation varies to the long antennae (*sa*^WT^) moth. This phenotypical difference appears in future generations, even at low temperature. Reversion is rare, and at low temperature the offspring of reverted individuals will continue the *sa* mutation (short antennae).

Many studies involved in epigenetic phenomenon concern DNA methylation. The data about the effect of diet on gene methylation and the release of hidden genetic variation by impairment of heat shock protein 90-mediated buffering systems offer eloquent examples of how epigenetic mechanisms might affect gene-environment interactions (Vercelli et al., 2004). But the change of methylation to demethylation is usually due to transition through the male gamete (Ariel et al., 1995, Biliya and Bulla, 2010).

Even though inheritance is possible along both the paternal and maternal lines, we focused on the analysis of inheritance along the paternal line. The paternal inheritance is more easier to study (Pavelka and Koudelová, 2001) because preparation of sperms is more simple than non fertilized eggs. In the present study, we analyzed the effects of alien sperms. Our epigenetic trait was also tranfered via sperm. We suppose that the long lasting epigenetic effect observed in our experiment evidence a simpler regulatory mechanism in insect cell, as compared to mammals (see Garcia-Fernández and Holland 1994, Morris et al., 2008).

Studies in mammals documents a different epigenetic phenomenon (Rassoulzadegan et al., 2006, Cuzin and Rassoulzadegan, 2010) that shares a considerable similarity with the phenomenon observed by us (Pavelka and Koudelová, 2001), especially in transmission of characteristics to subsequent generations. However, the phenomenon described by us is special due to the fact that it has been induced by a change in temperature during the development of an individual. Rassoulzadegan et al. (2006) report a modification of the mouse *Kit* gene in the progeny of heterozygotes with the null mutant *Kit*^tm1Alf^ (a *lacZ* insertion). Wild male mice mated with female mutant (heterozygous mouse *Kit^tm1Alf^*) have offspring with homozygous wild genes and white spots like the mutant female, even though they have no allele for these spots.

In spite of a homozygous wild-type genotype, their offspring maintain, to a variable extent, the white spots characteristic of *Kit* mutant animals (Rassoulzadegan et al., 2006). Large amount of a aberrant RNA is produced from the mutant gene (*Kit*^tm1Alf^) and accumulates in sperm and the defective RNA is transmitted to the embryo. Presence of the aberrant RNA silences the activity of the Kit wild-type gene then so the animals have white spots, even if they do not carry the corresponding mutant gene (see Soloway 2006). Epigenetic information is frequently erased at the start of each new generation. However, in some cases epigenetic information can be transmitted from parent to progeny (multigenerational epigenetic inheritance). A particularly notable example of this type of epigenetic inheritance is double-stranded RNA-mediated gene silencing in *Caenorhabditis elegans.* This RNA-mediated interference (RNAi) can be inherited for more than five generations (Buckley et al., 2012). Although the effect of piRNAs in *Caenorhabditis elegans* can survive to at least 20 generations (Ashe et al., 2012).

To understand this process, here we conduct a genetic screen for nematodes defective in transmitting RNAi silencing signals to future generations (Buckley et al., 2012). Between-generations epigenetic inheritance is not rare, for example Jablonka and Raz (2009) found over 100 cases of epigenetic inheritance in 42 species (bacteria, protists, plants, animals). However, the number of generations with epigenetic characteristics (epigenetic memory) are usually restricted. For example, in mice miRNAs the effect disappears after 6 generations (Rassoulzadegan et al., 2006). In grape phylloxera *Daktulosphaira vitifoliae* was epigenetic memory observed for 15 generations (Forneck et al., 2001), but in this case it was parthenogenetic transfer. It appears that small RNAs do not function without assistance. Buckley (et al., 2012) establish that the Argonaute protein HRDE-1 directs gene-silencing events in germ-cell nuclei that drive multigenerational RNAi inheritance and promote immortality of the germ-cell lineage.

We tested the stability of our epigenetic phenomenon and the frequency of the return to the original phenotype, because this component of epigenetic phenomena deserve more detailed study. Our phenomenon survives multiple generations (20 to date), though the descendants of one pair did revert to the original phenotype.

In this work we demonstrate that environmental influences including temperature (Pavelka and Koudelová, 2001), diet and dietary salts, give rise to epigenetic effect. We demonstrate transmission to future generations via RNA. In this study we focus on the cause of particular epigenetic phenomenon in insects, and their persistence through multiple generations.

## Materials and Methods

### Flour Moth Rearing and Handling

#### Animals and breeding

For experiments, the strain of the Mediterranean flour moth (*Ephestia kuehniella* Zeller, Lepidoptera: Pyralidae) homozygous for the autosomal recessive mutation *short antennae* (*sa*) was used. The strain was derived from a mutant Qy strain that was obtained from the stock cultures of W. B. Cotter, Jr. (Albert B. Chandler Medical Center, University of Kentucky, Lexington, KY) and has been kept in single-pair cultures at the Institute of Entomology (České Budějovice, Czech Republic) since 1991.

Stock cultures were reared in constant temperature rooms (20°C ± 1°C) at a 12-h: 12-h light/dark regime, and at a humidity level of about 40 %. Experimental and control cultures were kept at either 20°C ± 1°C at the same conditions. New generations were done with single-pair cultures. Pairs were collected during copulation and placed individually into empty Petri dishes (6 cm in diameter). Females laid eggs for 3-4 days, then imagoes were removed and Petri dishes with eggs were put into plastic boxes with food. Hatching larvae migrated from the dish to the food where they completed their development. Larvae were fed with milled wheat grains supplemented with a small amount of dried yeast.

#### Procedure

In order to determine the epigenetic effect we utilised multiple paramutant animals, kept in normal breeding conditions. We observed the ratio of short and long antennae in every generation. The exact number of animals with a specific phenotype were observed to F12. The ratio of phenotypes was relative constant, and other generations were simply observed visually without a complete count being made.

In this study, only short antennae (*sa*) and prolonged antennae (*sa*^WT^) with changed phenotype were distinguished. The first generation (*sa*^WT^ phenotype) of the line served to preparation of the changed phenotype (*sa* mutation). This mutation was turned out at 25°C ± 1°C, subsequent generations with the wild-type phenotype were then nursed at 20°C to exclude additional influence of temperature, and to monitor epigenetic feature only. The *sa* males of the wild phenotype (*sa*^WT^) were used for all subsequent generations. Each male of a *sa*^WT^ phenotype (*sa* genotype) was mated to a randomly chosen virgin female of the same phenotype and genotype.

In order to determine what is contained in the epigenetic information, the male spermatophore (from *sa*^WT^male) was analysed and its individual parts were then injected into the previously fertilized eggs. The eggs were from the original *sa* line, male and female *sa* phenotype, i.e. the line without epigenetic information.

The experimental design was governed by the method of exclusion: at first the impact of separate ejaculate components (*sa*^WT^ males) was tested, then we focused on the sperm - first on fractionated proteins, then on all of the sperm with RNase, and finally total RNA from the sperm *sa*^WT^ males was used. Similarly, eggs were injected with a solution of geldanamycin.

### Isolation of separate parts of spermatophore

Immediately after the copulation ended (*sa*^WT^male and female), the female was dissected. The following components from male spermatophore were separated for injection: (1) the spermatophore sac from the bursa copulatrix, (2) the seminal fluid from the spermatophore containing both the eupyrene and apyrene sperm, and (3) the secretion of male accessory glands from the bursa copulatrix. Ten samples of each category were stored at –70°C, and later homogenized and mixed with injection buffer (5 mM KCl, 0.1 mM NaH_2_PO_4_, pH 6.8) (Rubin and Spradling, 1982). Separated parts of spermatophore were prepared in room temperature only sperm into which RNase was added after homogenization (inactivated after 3 hours by incubating at 95°C for 30 seconds) were used for injections. Sperm proteins were purificated using Sep-Pak^®^ (Cartridges for solid Phase Extraction) Waters Corporation according to the manufacturer’s instructions. The acquired components were mixed together with injection buffer (Rubin and Spradling, 1982) and individual dissolutions were divided into several aliquots and deep frozen at –70°C. These aliquots have been used successively to inject fertilized eggs. Total RNA was extracted from sperm using the RNA Blue reagent according to the manufacturer's instructions (Top-Bio, Czech Republic). Inactivation test of some heat shock proteins was done by geldanamycin solution that is able to block these proteins. All chemicals were supplied by Sigma-Aldrich (Sigma-Aldrich s.r.o. Prague, Pobrezni 46 Czech Republic).

### Injecting Eggs

Total RNA, homogenized protein or geldanamycin were dissolved in injection buffer and approximately 0.2 -1 fmol of RNA, 0.05 μg proteins or 0.μg geldanamycin (ca 0.3 μ1 of injection buffer) were injected into the ventral side of the posterior domain of *Ephestia* embryos, similar to *Drosophila* or *Chymomyza* (see Pavelka et al., 2003). One-tree day old fertilized eggs were injected (Tab 1A).

**Tab. 1A.** Number of repetitions of each experiment and total number of imagoes used in experiment with additives on eggs.

| Experiment no. | additives injected to eggs | Numbers of repetitions | Total numbers of imagoes |
| --- | --- | --- | --- |
| 1 | *sperm* | 17 | 636 |
| 2 | *acc.glands* | 6 | 187 |
| 3 | *sperm.sac* | 5 | 152 |
| 4 | *pur.sperm* | 7 | 456 |
| 5 | *sperm+RNase* | 10 | 164 |
| 6 | *tot.RNA* | 7 | 140 |
| 7 | *geldan* | 11 | 353 |
| 8 | *buffer* | 7 | 390 |

### The environmental effect of food

Larvae (*sa*) were fed with milled wheat grains supplemented with a small amount of dried yeast with added NaCl or LiCl to 1.1 on 1mg of food. In another case, the larvae were fed only plain flour (Tab 1B).

**Tab. 1B.**
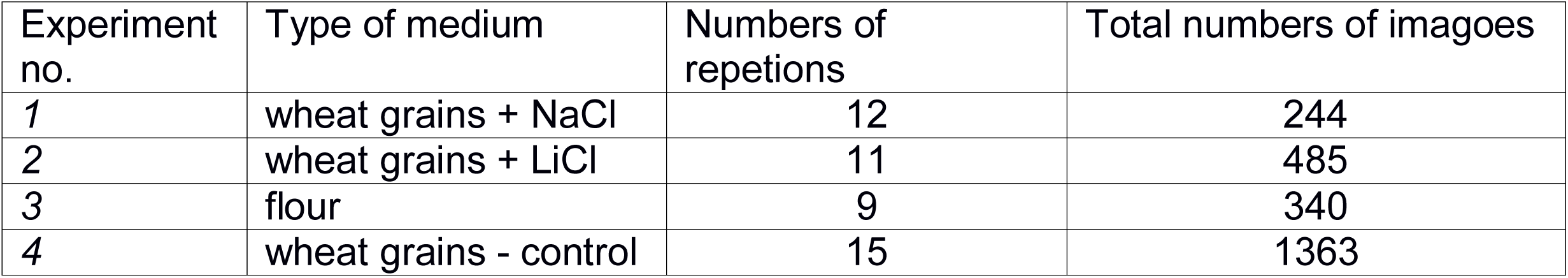
Number of repetitions of each experiment and total number of imagoes used in experiment with type of medium used for evaluating environmental effect of food.

### Monitoring of following generations

Over 40 lines carrying epigenetic information for twenty generations were monitored. We counted a proportion of epigenetic trait (*sa*^WT^) and nonepigenetic trait (*sa*) for individuals in generation (Tab 1C). F_6_ – F_11_ and F_13_ – F_20_ generations were just observed without precise counting.

**Tab. 1C.** Number of repetitions of monitoring of changes in proportion of epigenetic trait (*sa*^WT^) and nonepigenetic trait (sa) in following generations.

| Generation | Numbers of repetitions | Total numbers of imagoes |
| --- | --- | --- |
| *F1* | 43 | 4276 |
| *F2* | 54 | 4687 |
| *F3* | 53 | 4592 |
| *F4* | 42 | 4652 |
| *F5* | 42 | 4303 |
| *F12* | 44 | 5987 |

### Statistical Analyses

A number of repetition and number of imagoes are listed in Tab. 1. We counted percentage of *sa*^WT^ offspring of each moth´s pair and then we performed arcsin-transformation to meet demands of parametric statistical test – one-way ANOVA. We compared transformed percentage among all additives on eggs (injection of the sperms, the spermatophore sac, the secretion of male accessory glands, geldanamycin, clean proteins from sperms, sperms with RNase, RNA from sperms) or type of medium used for evaluating environmental effect of food (flour, wheat grains with NaCl, wheat grains with LiCl). Post-hoc comparison between every group was done by Tukey test for unequal N (Tab 2).

**Tab. 2.** The comparison between individual additives to the eggs, post-hoc Tukey test. The numbers in the table means statistical difference between the dyad of experimental groups. *sperm* – denaturated sperm, *acc*. *glands* - accessory glands, *sperm*. *sac* - spermatophore sac, *pur*. *sperm* - denaturated sperm purificated Sephadex, *sperm+RNase sperm* - denaturated sperm with Ribonuclease, *tot*. *RNA* – total RNA from *sa*^WT^ moth, *geldan* - injection antibioticum geldanamycin.

| additive | *sperm* | *acc.glands* | *sperm.sac* | *pur.sperm* | *sperm+RNA* | *tot.RNA* | *geldan* |
| --- | --- | --- | --- | --- | --- | --- | --- |
| *sperm* |  |  |  |  |  |  |  |
| *acc.glands* | 0.3 |  |  |  |  |  |  |
| *sperm.sac* | 0.1 | 0.9 |  |  |  |  |  |
| *pur.sperm* | 0.9 | 0.8 | 0.6 |  |  |  |  |
| *sperm+RNase* | 0.2 | 0.9 | 0.9 | 0.9 |  |  |  |
| *tot.RNA* | 0.9 | 0.13 | 0,057 | 0.8 | 0,01 |  |  |
| *geldan* | 0.003 | <0.001 | <0.001 | 0.004 | <0.001 | 0.3 |  |
| *buffer* | 0.6 | 0.9 | 0.9 | 0.9 | 0.9 | 0.3 | <0.001 |

Monitoring of changes in proportion of epigenetic trait (*sa*^WT^) and nonepigenetic trait (*sa*) in following generations was tested by contingency tables. Non-outlier maximum of the dataset is 12,6 % of reverted individuals in a clutch. For the statistical tests, we consider clutches that have more then 12,6 % of reverted individuals as reversed, less than the percentage as normal. We compared generations F1, F2, F3, F5 and F12.

All tests were performed in program R 3.3 (2016 The R Foundation for Statistical Computing).

## Results

### Injections

There are differences in percentage of *sa*^WT^ offsprings between different types of additive to the eggs (ANOVA, F = 9,49, df = 7, p < 0.001, post-hoc tests in Tab. 2) (Fig. 1). Injection of *sa*^WT^ denaturated sperm to fertilise eggs (*sa* x *sa*) did not change the proportion of long antennae in offspring. The same null effect was observed when injected with extracts from accessory glands, spermatophore sac, purificated sperm, and sperm with Ribonuclease (RNase) or geldanamycin. However, the injection of total RNA isolated from *sa*^WT^ sperm, into fertilized eggs (*sa* x *sa)* produced a significantly higher percentage of long-antennae offspring.

**Fig 1.**
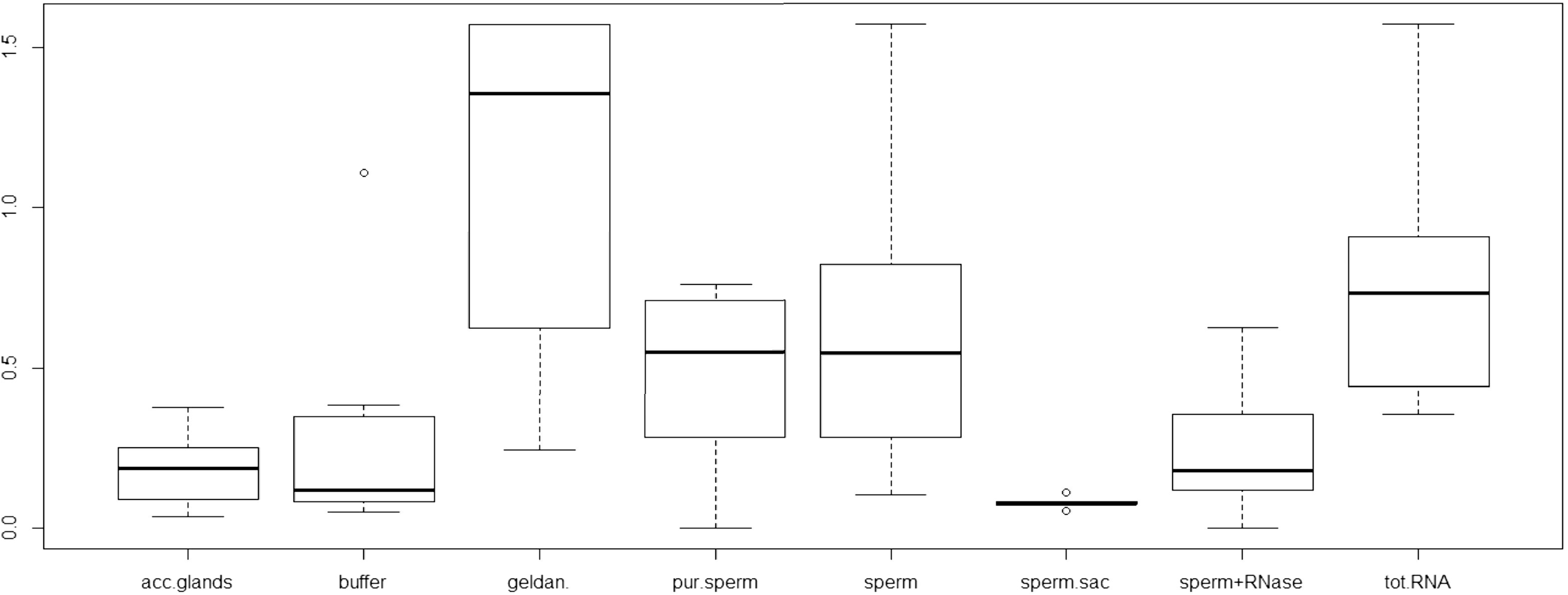
The differences in percentage of *sa*^WT^ offsprings between different types of additive to the eggs (shortcuts see Tab 2). The central thick line is median, longer thin ones upper and lower quartiles, short thin ones are non-outlier maximum and minimum, circles are outliers.

### Epigenetic memory

During the monitoring of generations of one-pair cultures, we found out that there is a reversion to beginning mutant phenotype by some individuals in every generation (Fig. 2). Number of reverted individuals was usually 0 – 6% per generation and pair. Rarely, the offspring of some pairs reverted in large number (cca 90 %). We have tried to determine the point at which there is a revertion of a significant number of individuals in populations. We analysed number of extremely reversed clutches counted as percentage of reversed individuals higher than non-outlier maximum for whole dataset which is 12,6. There is no change in proportion of these extreme clutches between generations F1, F2, F3, F5 and F12 (chi-square = 2,6, df = 4, p = 0,6).

**Fig 2.**
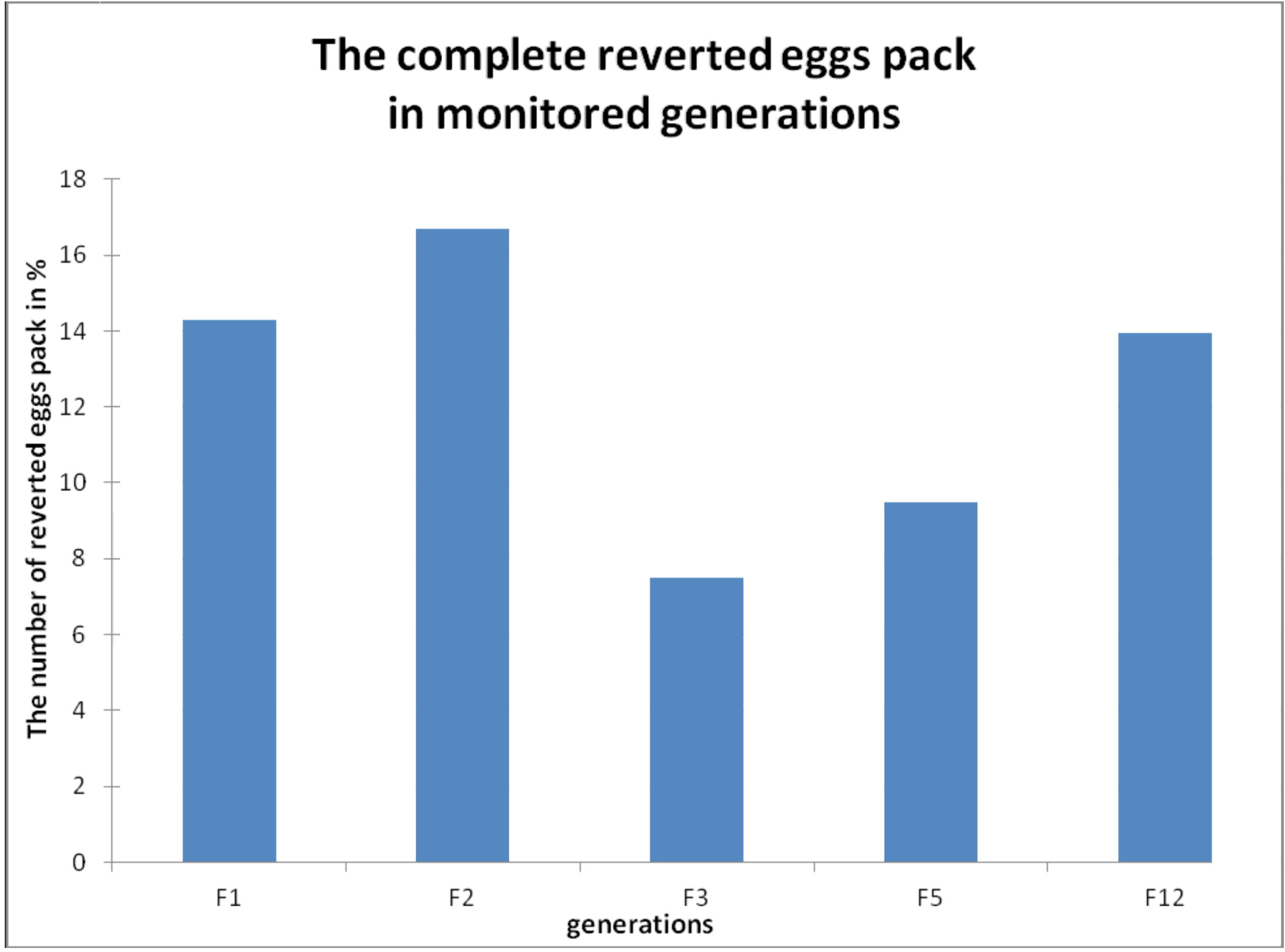
Number completely reverted clutches precisely evaluated in generations.

Tracking progeny was conducted for each generation, but the precise evaluation of all specimens was carried out for generations F2, F3, F5 and F12. For other generations, including F20, the ratios of reverted individuals were evaluated, as with similar cases (the calculation was not carried out in all cultures). Within 5 control cultures the epigenetic effect continued to the 40th generation, and then terminated.

### The effect of salts and poor nutrition

The percentage of phenotype changes (*sa* on *sa*^WT^) differed by diet (F = 5,76, df = 2, p = 0,005). Clutches fed with NaCl had a lower proportion of phenotypically altered individuals, then those fed without added salts (p = 0,005). The other pair combination show no differences (Fig 3).

**Fig 3.**
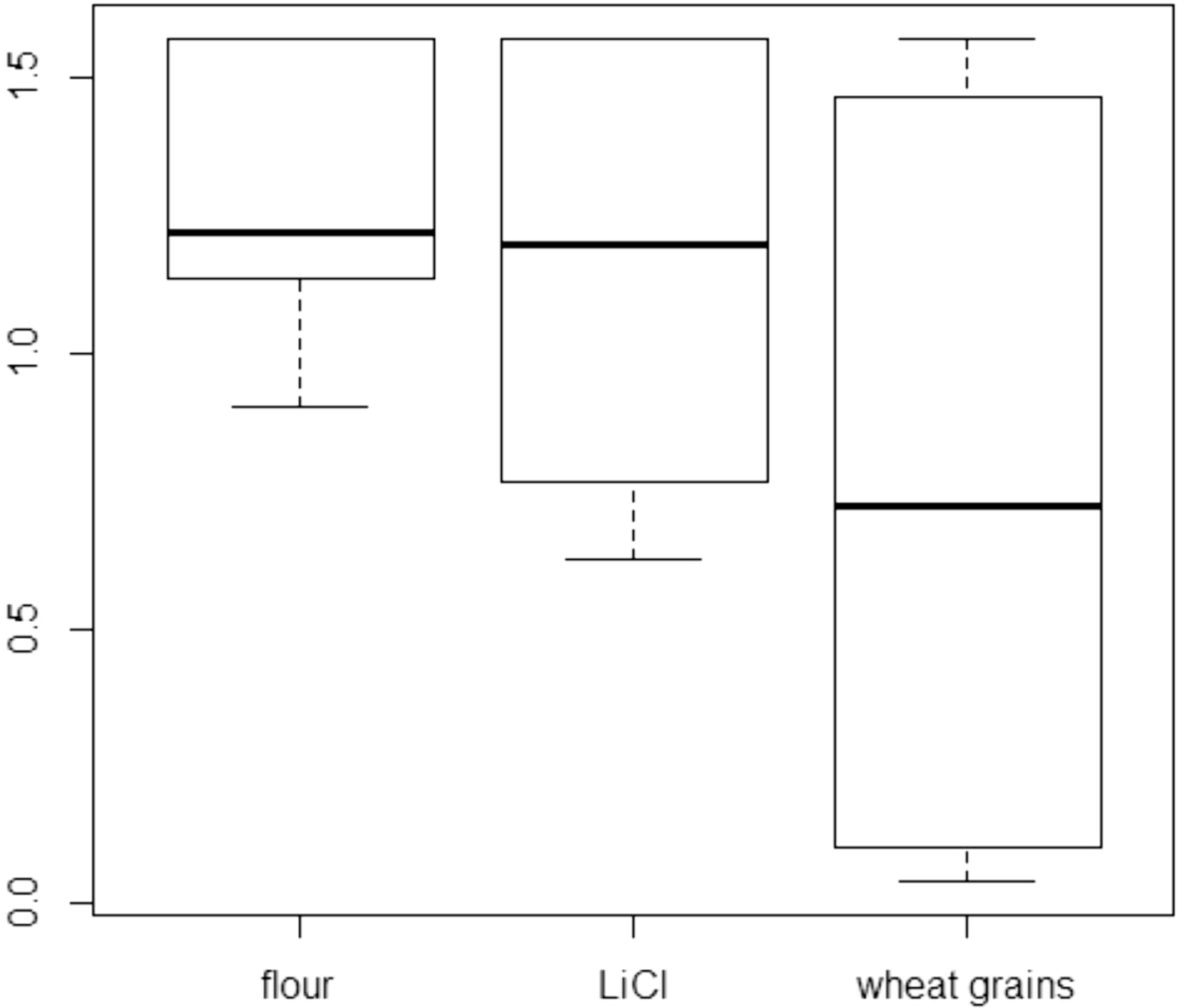
The differences in percentage of *sa*^WT^ offsprings between different types of food. The central thick line is median, longer thin ones upper and lower quartiles, short thin ones are non-outlier maximum and minimum.

## Discussion

Results of our experiments on lepidopterans revealed that epigenetic effect was mediated by RNA and lasted for 20 (or 40) generations. It is possible that epigenetic effect would continue further to next generations. It is probably the longest duration of epigenetic influence ever observed in insect. We demonstrated significant infuence of total RNA isolated from sperm of males on changed phenotype of antennae caused by epigenetic effect. The other components (accessory glands, spermatophore sac, denaturated sperm with Ribonuclease) separated from sperm of males carrying epigenetic effect did not have analogous impact. The influence of geldanamycin comparable RNA injection was even stronger than expected, confirming that the effect is due to the small RNA. Geldanamycin inhibits heat shock protein Hsp90 (Whitesell et al., 1994). Heat shock protein 90 (Hsp90) is a molecular chaperone essential for activating many signaling proteins in the eukaryotic cell (Pearl and Prodromou, 2006). The link between small RNA and Hsp90 heat shock protein and Argonaute proteins (Argonaute proteins bind different classes of small RNAs) has been demonstrated (Iwasaki et al., 2010).

A similar effect of temperature was observed in *Caenorhabditis elegans*. A majority of the SynMuv B mutants grown at high temperature were irreversibly arrested at the L1 stage. High temperature arrest (HTA) is accompanied by upregulation of many genes characteristic of the germ line, including genes encoding components of the synaptonemal complex and other meiosis proteins (Petrella et al., 2011). Unfortunately the link to small RNAs is unclear.

This phenomenon has been documented in mammals (see Rassoulzadegan et al., 2006). As the RNA is not degraded and continues to act in the cells, there might be a relationship with RNAs generating interference RNA (RNAi) (see Soloway, 2006). In this process also seems essential RNA methyltransferases. Research in animal models has shown that RNAs can be inherited and that RNA methyltransferases can be important for the transmission and expression of modified phenotypes in the next generation (Liebers et al., 2014, Kiani et al., 2013). Perhaps it increases the stability of small RNAs such as cytosine-5 methylation, are known to stabilize RNAs (Tuorto et al., 2012, Kiani et al., 2013).

Similar phenomenon conditioned by miRNAs found in lepidopterans has been observed for example in plants, where a temperature-dependent epigenetic memory from the time of embryo development expresses in norway spruce. This epigenetic machinery thereafter influences the timing bud phenology (Yakovlev et al., 2010).

Nevertheless, reversion of mutant phenotype occurs spontaneously also in the part, or rarely in all the offspring of studied flour moths (Fig.2). We suppose it means simple diminishing of epigenetic influence on the basis of gradual decrease of small RNAs (maybe piRNA) during ontogeny under particular treshold. So, an individual posses epigenetic trait, but all its tissues contain no small RNAs causing epigenetic effect any more. It could occur during ontogeny of some individuals, whereas others could retain the same concentration of small RNAs. Occasionally, in rare cases could some small RNAs disappear only in the parental tissues where eggs and sperm develop, while the individuals and their other tissues display epigenetic phenotype. The offspring then do not display epigenetic phenotype.

The epigenetic effect in flour moths is maintained even after many generations (our experimental culture has 20 monitored generations and total 40 observed). The piRNAs at *Caenorhabditis elegans* can trigger highly stable long-term silencing lasting at least 20 generations. Once established, this long-term memory becomes independent of the piRNA trigger but remains dependent on the nuclear RNAi/chromatin pathway (Ashe et al., 2012). Insects benefits more from the pronounced variability of the progeny, because they produce much larger numbers of offspring and are physiologically more influenced by environmental conditions than mammals.

Epigenetic memory in mice lasts only six generations (Rassoulzadegan et al., 2006), in voles is the memory documented just to F3 generation (Francis et al., 1999). However, the situation is different in insects. The studied epigenetic trait (developed antennae) is involved in finding of sexual partner (see Pavelka and Koudelová, 2001) and lasts for many generations. It was not observed in mammals. Such long lasting epigenetic effect could be considered potentially adaptive (Forneck et al., 2001; see Loxdale, 2010). It is possible that epigenetic phenomena could have evolutionary consequences that increase variability in offspring.

Many studies have demonstrated that epigenetic mechanisms, including DNA methylation and histone modification, not only regulate the expression of protein-encoding genes, but also miRNAs (Sato et al 2011). Conversely, another subset of miRNAs controls the expression of important epigenetic regulators, including DNA methyltransferases, histone deacetylases and polycomb group genes. This complicated network of feedback between miRNAs and epigenetic pathways appears to form an epigenetics-miRNA regulatory circuit, and to organize the whole gene expression profile (Sato et al., 2011, Wei et al., 2010).

Our results show that the phenomenon persists for many generations (beyond the 20 monitored). A number of individuals revert to the genetically conditional mutant phenotype. Generations for which accurate images existed were processed statistically. There is no change in proportion of these extreme clutches between generations. The number of reverted individuals is 10-15%. The studied epigenetic phenomenon effects about 85-90% of the population. The population variability is ensured This effect is therefore evolutionarily significant. It is possible that switching between states occurs because certain small RNA in the fertilized egg are lower than the critical concentration required. However, the mechanism can be far more complex.

The most common manifestation of epigenetic change it can be seen in the sirtuin genes. The similarity with our phenomenon is primarily in the long-term inheritance genotype, which is occasionally interrupted for reasons which remain unclear. But Sirtuin genes also cause the *sa* phenotype *that result from heat, starvation and salts*.

White–opaque switching in the human fungal pathogen *Candida Albicans*, results from the alternation between two distinct types of cells (Sonneborn et al., 1999, Zordan et al., 2006). Switching is probably caused by the SIR2 (silent information regulator) gene, which seems to be important for phenotypic switching (Pérez-Martín et al.,1999). Silent Information Regulator (SIR) proteins are involved in regulating gene expression and some SIR family members are conserved from yeast to humans (Sherman et al., 1999). SIR proteins organize heterochromatin at telomeres (Palladino et al., 1983). SIR genes have many functions. Sirtuins are evolutionary conserved NAD(+)-dependent acetyl-lysine deacetylases and ADP ribosyltransferases dual-function enzymes involved in the regulation of metabolism and lifespan (Balcerczyk and Pirola, 2010). Sirtuins are to hypothesized to play a key role in an organism's response to stresses (such as heat or starvation). A calorie restriction turns on a gene called PNC1, which produces an enzyme that rids cells of nicotinamide, a small molecule similar to vitamin B3 that normally represses Sir2. The gen PNC1 is also stimulated by other mild stressors known to extend yeast life span, such as increased temperature or excessive amounts of salt (Sinclair and Guarente, 2006).

A similar effect is observed in the epigenetic sa mutation, resulting from the stress of bad nutrition, the effect of increased temperatures, and the presence of salts.

## Related Sirtuin genes and small RNA?

The characteristics of this phenomenon may be related to SIR proteins, though secure transfer has only been experimentally demonstrated through RNA. We believe that SIR proteins in addition to various known functions (e.g. silence genes - see Ralser et al 2012) might also catalyze the formation of small double-stranded RNA, according to damaged genes to be silenced. SIRT regulation is multifaceted, but not yet considered to be associated with RNA. We present the hypothesis that insects can initiate the creation of an RNA against harmful genes. Deleterious gene, form their RNA SIRT is apparently identifies and starts producing a Piwi RNA that maintains the epigenetic regulation, or to interact with SIRT system. Transmission via small RNAs is proven, and we do not expect any impact on the epigenetic methylation phenomenon. In the reference case, the epigenetic influence is caused by an RNA, and probably one of the categories of small RNAs. Small regulatory RNAs such as short interfering RNAs (siRNAs), microRNAs (miRNAs), and Piwi interacting RNAs (piRNAs) have key regulatory functions (Peters and Meister, 2007).

SIR genes linked with small RNAs have been reported. It was shown that small interfering RNA-mediated SIRT7 knockdown leads to reduced levels of RNA Pol I protein, but not messenger RNA, which was confirmed in diverse cell types (Tsai et al., 2012). Regarding small RNAs studied, the mechanism best suits piRNAs transmission, through transmission via sperm, for example. The piRNA exists in invertebrates and mammals and piRNAs are abundant in testes (Aravin et al. 2006). However, it is noted that the majority of piRNAs are antisense to transposon sequences (Malone and Hannon, 2009, Webster, et al., 2015). Drosophila Piwi-family proteins Aubergine (Aub) a Argonaute-3 (Ago3) have been implicated in transposon control (Brennecke et al., 2007, Li et al., 2009, Zhang et al., 2011). Gene silencing may be carried out by transposons or, as in our study, by some other small RNA with a similar function.In the case of the sa mutation small RNAs neutralize harmful mutation. Males with this mutation have difficulty finding mates (Pavelka and Koudelová, 2001). It is possible that Piwi system and tertiary siRNAs may neutralize other harmful mutations, and somehow connected with sirtuin genes. Apparently, as with *E. kuehniella*, some important regulatory genes are activated epigenetically. The observed results correspond to the effect of sirtuins genes and piRNA. We have a hypothesis that these mechanisms are closely linked in some way, even if we are not able to determine how.

In summary, the effect of small RNAs in many insect generations was observed for the first time. We mapped reversion of mutations to the wild phenotype, though it is not clear why this usually occurred only in a small proportion of offspring and rarely on the whole population. Our study demonstrated that epigenetic factors appear to effect phenotype by blocking factor as heat shock proteins. As a result, it is thought that SIRT genes could be regulated by small RNAs.

## Acknowledgments

The research and findings reported in this study were conducted and delivered before funding and research support was terminated in 2004.Thanks to “help” František Marec and Vladimír Košťál this manuscript had not been submitted (published) at time when it could be completely ground-breaking. At present, can only sort among many similar manuscripts.

